# Evolution of mating types in finite populations: the precarious advantage of being rare

**DOI:** 10.1101/400598

**Authors:** Peter Czuppon, David W. Rogers

## Abstract

Sexually reproducing populations with self-incompatibility bear the cost of limiting potential mates to individuals of a different type. Rare mating types escape this cost since they are unlikely to encounter incompatible partners, leading to the deterministic prediction of continuous invasion by new mutants and an ever increasing number of types. However, rare types are also at an increased risk of being lost by random drift. Calculating the number of mating types that a population can maintain requires consideration of both the deterministic advantages and the stochastic risks. By comparing the relative importance of selection and drift, we show that a population of size *N* can maintain a maximum of approximately *N*^1/3^ mating types for intermediate population sizes while for large *N* we derive a formal estimate. Although the number of mating types in a population is quite stable, the rare type advantage promotes turnover of types. We derive explicit formulas for both the invasion and turnover probabilities in finite populations.

## 1. Introduction

Splitting a population of universally compatible gametes into two or more self-incompatible mating types can potentially limit inbreeding depression, control the spread of selfish organelles, and help mate searching (Hoekstra, 1982; Hurst and Hamilton, 1992; Charlesworth et al., 2005; Billiard et al., 2011; Hadjivasiliou and Pomiankowski, 2016). Although the origins and exact benefits of producing different mating types remain controversial (Perrin, 2012), the costs are clear: self-incompatibility reduces the number of potential mating partners and increases the potential for subfertility (Fisher, 1930; Power, 1976). However, if gametes are compatible with any class but their own, new (and therefore extremely rare) mating types do not pay this cost: they can mate with any member of the population. Hence, novel mating type mutants should increase in frequency until all types are equally represented. The deterministic outcome of mating type evolution is the inexorable rise in the number of different types driven by negative-frequency dependent selection for rare mutants, also referred to as the “rare sex advantage” (Iwasa and Sasaki, 1987). Few studies have considered the flip-side of the rare sex advantage in the context of haploid self-incompatible mating types, i.e. rarer types are at much greater risk of being lost from finite populations by random genetic drift (Douglas et al., 2016; Constable and Kokko, 2018). These stochastic extinction events not only hinder invasion of rare mating types, they also prevent the population from maintaining too many low-frequency mating types.

Multiple different genetic mechanisms of mating type determination have evolved, including those regulated by two unlinked loci like the tetrapolar mating systems of many fungi hypothetically capable of generating hundreds of compatible mating types, and the more widespread single locus incompatibility systems. Here, we focus on a system of mating type determination typified by the yeast *Saccharomyces cerevisiae* where a single haploid locus is responsible. Despite the very strong negative frequency-dependent selection for rare mating types, most single locus haploid incompatibility systems are binary. They produce only two mating types, similar to the two sexes found in many animals, despite a population with only two types experiencing the highest rate of incompatible matings (Hurst, 1996). Even in non-binary systems, only a few additional mating types are typically present (James, 2015). For example, heterothallic populations of slime molds, including the social amoeba *Dictyostelium discoideum*, can contain 2-13 different mating types (Bloomfield et al., 2010; Clark and Haskins, 2010; Douglas et al., 2016). Similar numbers, 2-12 different mating types have been observed in ciliates for the *Tetrahymena*-species (Eduardo and James, 1964; Doerder et al., 1995; Phadke and Zufall, 2009). A high diversity of mating types has been reported for certain mushroom-forming fungi: the global population of fairy inkcap *Coprinellus disseminatus* is estimated to have 123 mating types, but the number found in a single population would inevitably be considerably smaller (James et al., 2006). There is an upper limit to the number of different mating types that can be maintained by a finite population – a number low enough for the strength of negative frequency-dependent selection to not be overwhelmed by genetic drift. Here, we set out to determine where this threshold lies.

We present a model of haploid self-incompatibility that allows estimation of the number of mating types in a finite population by comparing the deterministic and stochastic dynamics of mating type frequencies. The simplicity of this heuristic approach, i.e. a non-rigorous analysis under simplifying assumptions, allows an analytical solution, generating a straightforward estimation of the maximum number of mating types that can be maintained in a small population. Furthermore, this approach allows derivation of the probabilities of invasion of a rare mating type and the turnover of mating types. This method relies on the assumption that large stochastic deviations from the stationary distribution result in extinction, an assumption which becomes less tenable when rare mating types have larger absolute representation. To evaluate the accuracy of our heuristic approach, we compare it to a rigorous derivation of extinction probabilities in the spirit of (Wright, 1939) which cannot be solved analytically for our model. We show that our approximate analysis agrees with the numerical evaluation arising from the rigorous approach for small to intermediately sized populations. Our approach is similar to a recent analysis (Constable and Kokko, 2018) where an estimate for the upper and lower bounds for the number of mating types was derived, except that instead of building on the estimation of the stationary distribution on the level of mating types, we are able to provide a precise numerical estimate by building on the estimation of the stationary distribution of a focal mating type.

## 2. Methods

### 2.1 Model

In order to quantify the effects of stochastic drift, mutation and negative frequency-dependence we consider a haploid population of *N* individuals in its demographic equilibrium, i.e. *N* is constant over time. Each individual is of a certain mating type, *M*_1_, …, *M*_*R*_ for some positive integer *R*. The total number of individuals of type *M*_*i*_ is denoted by *X*_*i*_. Individuals of type *M*_*i*_ can only mate with individuals of a different type *M*_*j*_ ≠ *M*_*i*_. Hence, gametes are self-incompatible and can not reproduce asexually.

We implement changes in the population configuration by the Moran process, generations are overlapping. A transition in the population configuration consists of a birth and a death event in order to maintain a constant population size. The order of events does not affect the overall dynamics and can be exchanged. This can be seen by reordering the terms in the transition rates below. In the birth-step two randomly chosen individuals have the opportunity to mate with each other. If the two individuals have different mating types they give birth to an offspring which randomly inherits the mating type of one of its parents. In case the two parents carry the same mating type, reproduction is not possible and the system remains in its current state. In the death-step a third randomly chosen individual gets replaced by the newly born offspring thus leaving the population size unchanged. Note that instead of drawing a third individual here, we could also choose an arbitrary individual of the whole population including the parents. Since we will work with intermediate to large population size approximations, this does not alter the results. Also, the dynamics remain the same except for the case of exactly two mating types where our implementation ensures that the population is always able to reproduce, thus producing a distinct boundary behavior. For an illustration of our haploid self-incompatibility system see also Figure 1.

**Figure 1:**
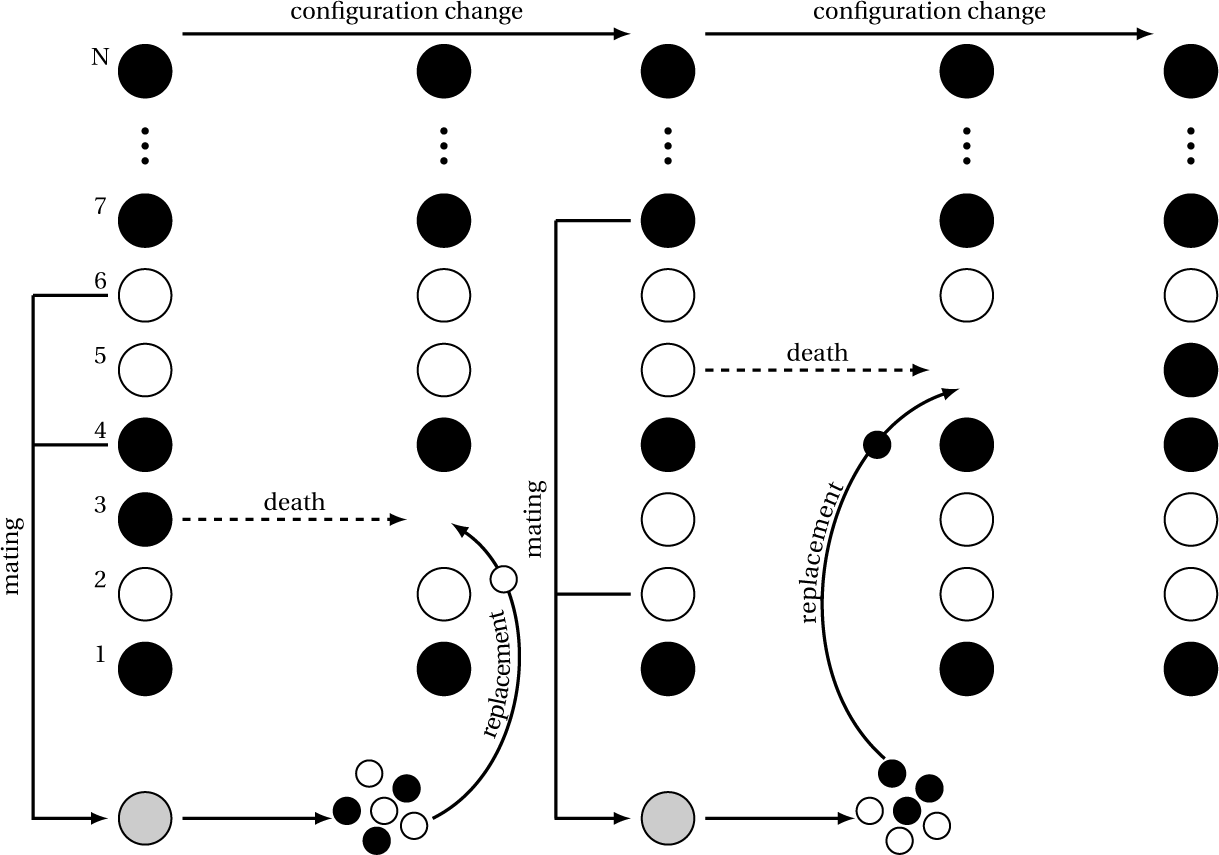
Mating between compatible individuals produces an offspring that replaces a randomly selected member of the population. In each time step three different individuals are chosen from the population. Two of them mate and produce a zygote which produces a large amount of gametes. One of these gametes is randomly chosen to replace the third initially drawn individual. The mating type of the offspring is chosen uniformly at random from the parental mating types. However, reproduction is only successful if the parents have different mating types, i.e. the gametes are self-incompatible.

The described dynamics translate into the following transition rates: the change from *k* to *k* + 1 individuals of mating type *M*_*i*_ happens at rate

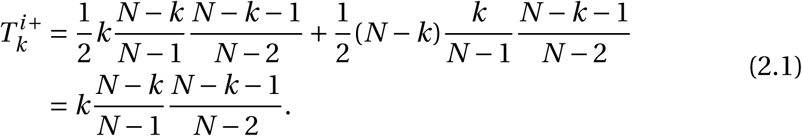

The single terms emerge as follows:

- The two terms in the first line arise due to reproduction events where either a *M*_*i*_-individual chooses a non-*M*_*i*_ individual or vice versa.
- The factor 1/2 is due to the offspring inheriting its mating type uniformly at random from one of the two parents. In half of the successful reproduction events including a *M*_*i*_-type parent it will inherit type *M*_*i*_.
- The *k*-term (or (*N* −*k*)-term) is the rate of drawing an individual of type *M*_*i*_ (or non-*M*_*i*_). The idea is that all of the *N* individuals have a random waiting time (exponential waiting time with rate 1) until they start the reproduction cycle which is independent of all the other individuals. Thus, having *k* type *M*_*i*_ individuals yields a rate *k*. This might seem to introduce a bias towards more frequent mating types but effectively does not since reproductive success is still coupled to finding a compatible mating partner.
- (*N* −*k*)/(*N* − 1) is the probability of drawing a non-*M*_*i*_ individual from the remaining *N* − 1 individuals.
- The last fraction (*N* −*k*−1)/(*N* −2) is the probability of a non-*M*_*i*_ individual dying and thus being replaced by the offspring. Note, that the parents are excluded from this set of individuals. This assures that there are always at least two mating types in the population.

Arguing analogously, we can write down the decrease rate, i.e. the rate to transition from *k* to *k* − 1 individuals of mating type *M*_*i*_:

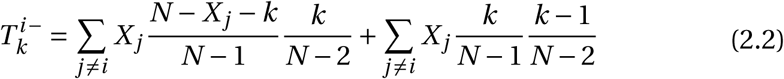

Again, we can disentangle the transition rate into single terms. The structure of the terms that are summed up is similar to the ones obtained for 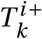. Further-more, the sums are interpreted as follows:

- The first sum describes matings which happen between non-*M*_*i*_ mating types resulting in a replacement of an *M*_*i*_ individual.
- The second sum is the rate at which a mating between a *M*_*j*_ and a *M*_*i*_ individual causes a *M*_*j*_ offspring to replace another *M*_*i*_ individual.

We note that instead of using transition rates we could also implement the model by transition probabilities per unit time step. Here we would replace the factor *k* by its frequency *k*/*N*. Another possibility is to condition on successful reproductive events to happen which would introduce a factor 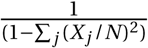.

Both of these alternative implementations would result in the same qualitative dynamical behavior, however on a different time-scale. Our implementation is motivated by aligning the time-scale of the individual based model with the corresponding deterministic system.

Using our defined transition rates, 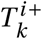 and 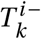, and assuming large population sizes *N* we can derive a stochastic differential equation (diffusion approximation – see the supplementary information (SI)) describing the dynamics of our model. Writing 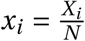 as the frequency of mating type *M*_*i*_ in the population, the stochastic differential equation reads as

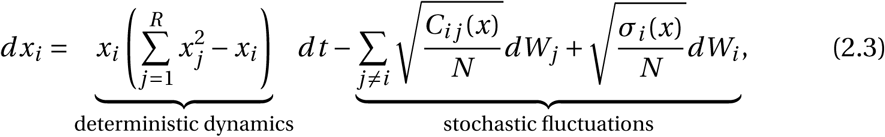

where (*W*_*i*_)_*i =*1,…,*R*_ are independent standard Brownian motions. The covariances between the focal mating type *i* and another mating type *j* are given by

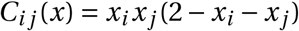

and the variance reads

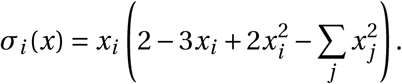

The stochastic diffusion (equation (2.3)) can be interpreted as follows: the first part describes the deterministic dynamics of the system which were already studied in (Iwasa and Sasaki, 1987) and more recently adapted in the framework of asymmetric mating choices in (Hadjivasiliou and Pomiankowski, 2016). The second term consisting of the sum over the non-*M*_*i*_ mating types describes the covariances which influence type-*M*_*i*_ individuals due to the restriction of a constant population size while the last term can be attributed to random births and deaths of type-*M*_*i*_ individuals, i.e. the dynamical development of the variance of *x*_*i*_. These make up the stochastic fluctuations of the finite size population, commonly referred to as genetic drift.

It is worth noting that using a Wright-Fisher implementation of the process, i.e. non-overlapping generations, yields the same dynamics (under an adequate time-rescaling), see for instance (Etheridge, 2012) on that issue. In the following we choose to use the Moran process since the analysis of invasion and turnover probabilities in Section 3.2 derives more naturally in this framework. The analysis of the mean number of mating types in Section 3.1 can be done analogously assuming Wright-Fisher dynamics.

In Figure 2 we show the temporal evolution of the system with four mating types in a population of (a) 100 and (b) 1, 000 individuals. The dashed lines represent the corresponding deterministic dynamics which all quickly collapse to the globally attractive equilibrium which is located at (1/4, 1/4, 1/4, 1/4), i.e. a uniform distribution of mating types. As can be seen, the stochastic trajectories remain closer to these lines in larger populations while smaller systems are more prone to random fluctuations. In Figure 2a eventually, due to a random fluctuation, one of the mating types goes extinct at which point the internal equilibrium shifts to 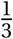 For a detailed study of the deterministic system we refer to the “Mating kinetics I” model in (Iwasa and Sasaki, 1987).

**Figure 2:**
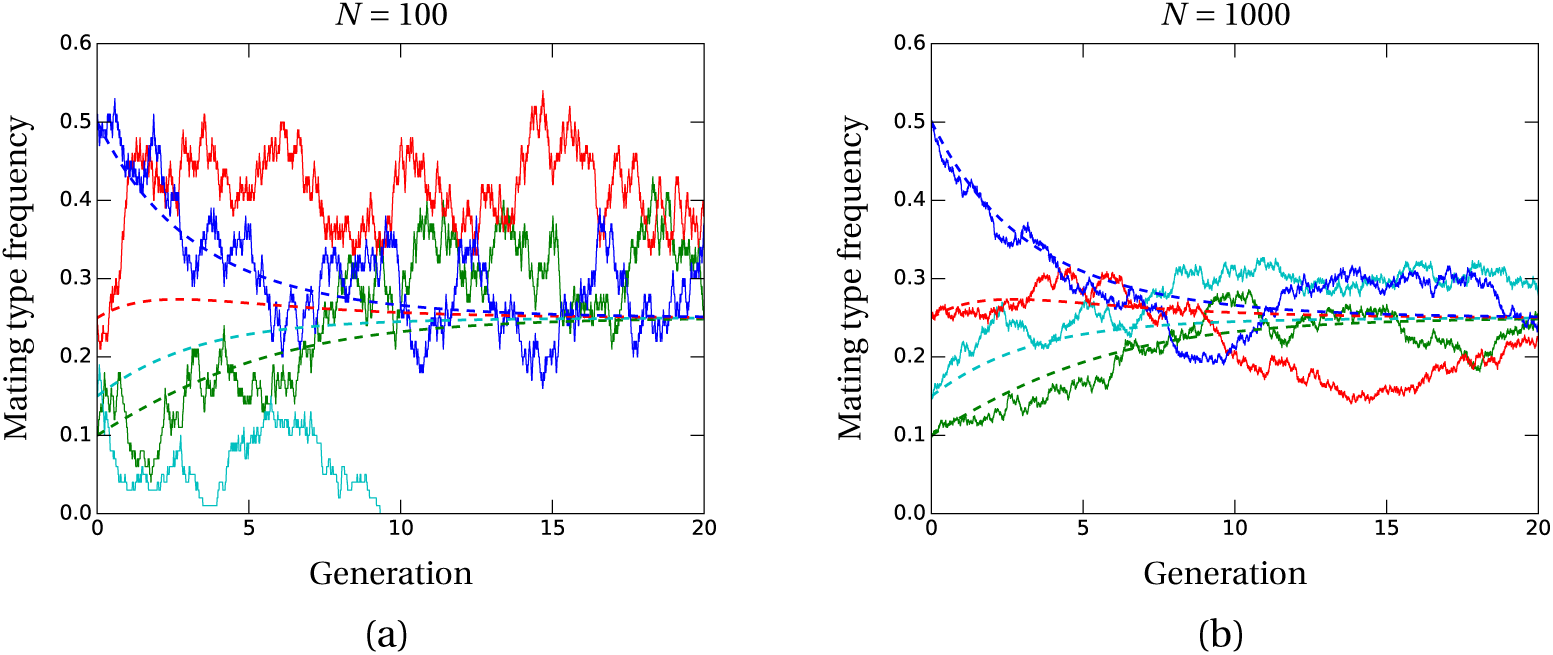
Mating types are driven to equal frequencies in large populations but more easily lost to genetic drift in small populations. Stochastic evolution of the number of mating types is plotted by solid lines in a population with (a) 100 and (b) 1000 individuals. While in small populations genetic drift overcomes the stabilizing deterministic effect, the contrary is true for larger population sizes. Dashed lines are deterministic (infinite population size limit) trajectories of the system.

### 2.2 Model including mutations

So far, we have described the dynamics of a population in which no new mating types emerge. Hence, the number of mating types is decreasing over time due to stochastic extinction events as expected in a finite population. These dynamics are illustrated in Figure 3a. Eventually, the number of mating types will collapse to two. Now, we proceed by randomly introducing new mating types (mutations) into the population. More formally, at every replacement step a new mating type (or mutant) emerges with probability *u*. The equations describing the model with mutations are stated in the SI. Since in the following we restrict ourselves to the case with low mutation rates, for the analysis it is sufficient to consider the previously derived formulas.

**Figure 3:**
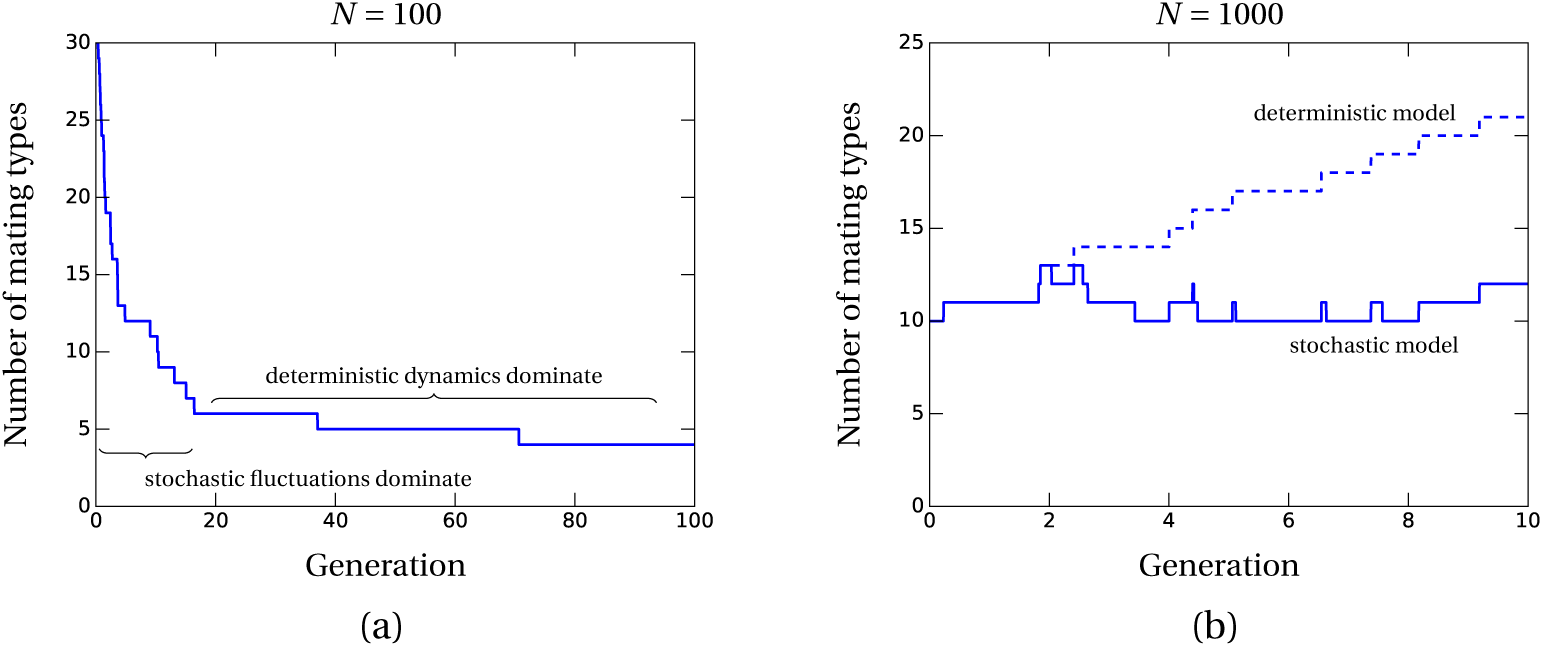
The balance between stochastic and deterministic effects determines the maximum number of mating types. (a) In a population of 100 individuals and initially 30 mating types genetic drift causes a lot of the initially present alleles to disappear within the first few generations. However, once a critical number of mating types is reached the stabilizing deterministic dynamics take over such that extinctions become rare. Subfigure (b) shows the evolution of the number of mating types under deterministic and stochastic dynamics in a system with mutations. While in the deterministic model (dashed line) the number of mating types is a monotonically increasing function over time, new mutants do not necessarily invade the population in the stochastic model (solid line). Further, due to extinctions the number of mating types can also decrease in the stochastic setup. The mutation rate is chosen as *u = N*^−1^ with *N =* 1000, meaning that on average there is one mutation per generation (*=* 1000 updates). The choice of this rate solely serves illustrative purposes in this plot. In subsequent figures it will be significantly reduced.

## 3. Results

### 3.1 Number of mating types in a finite population

The maximum number of mating types that a population can support depends on the balance between the deterministic and stochastic dynamics. In a finite population, the number of mating types *R* cannot exceed the total number of individuals *N*. Negative frequency-dependent selection will drive the frequency of each mating type towards the mixed equilibrium where all types are at equal frequency 1/*R*. The global stability of this equilibrium not only attracts the trajectories but also makes it unlikely, though not impossible, for mating types to die out by stochastic fluctuations once they are established.

The likelihood that a mating type is lost through stochastic fluctuations depends on the absolute number of individuals of that mating type in a population (Figure 3a) which becomes smaller for increasing numbers of mating types *R*; the stochastic dynamics are therefore responsible for the low number of mating types (small *R*) we expect to observe in small populations (small *N*). New mating types can appear through mutation. In a deterministic model, this results in an endless increase of the number of mating types over time (Figure 3b). However, in a finite population, adding a new mutant will increase the number of mating types in the population above the number where the risk of extinction is negligible. Eventually, a mating type will be lost by extinction and the system will return to the previous number of mating types where the stochastic and deterministic dynamics are balanced (selection-extinction balance). When mutants appear infrequently (small *u*), this balance is restored before another mutant appears and the mutation rate will have no effect on the predicted number of mating types. However, if the mutation rate exceeds the extinction time of the population with a certain number of mating types, then new mating types will accumulate in the population faster than they can be purged by stochastic fluctuations, and the observed number of mating types will increase up until a mutation-extinction balance is reached.

We show that in a finite population of size *N* with a low mutation rate we would expect a maximum of order *N*^1/3^ different mating types for intermediate values of *N* (Figure 4), see the derivation in SI, Section 2.1. That is, despite the selective advantage enjoyed by rare mutants – which depends on the resident number of mating types *R* as we will see later – a population of 1, 000 individuals could only support approximately 10 different mating types. We obtain this prediction by employing an order analysis of the dynamics of a focal mating type. We first calculate the highest order terms in the deterministic and stochastic parts of equation (2.3), and then determine the transition point where the deterministic dynamics are of higher order than the stochastic dynamics causing the population to stabilize.

**Figure 4:**
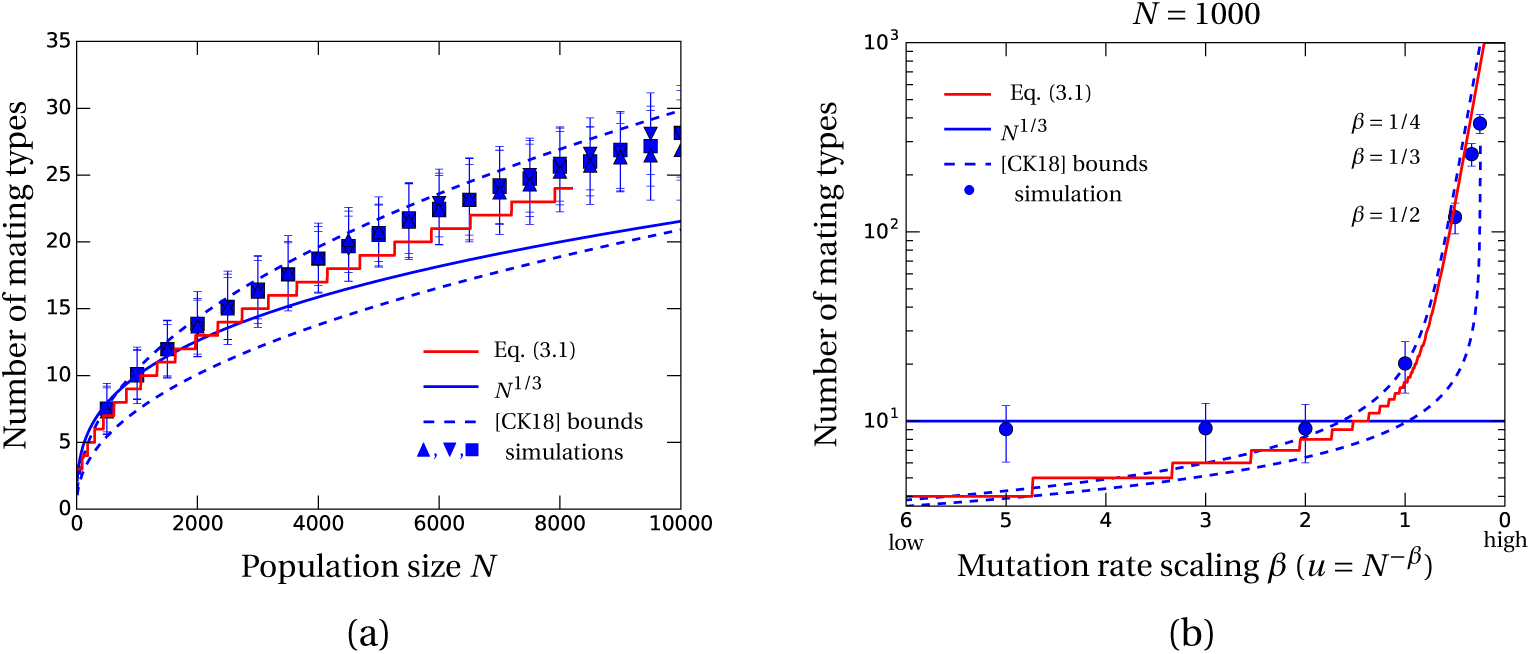
Simulations support the analytical solutions. (a) Comparison between the heuristic and rigorous prediction to simulations which started with 50 (down arrowheads), 20 (squares) or 3 (up arrowheads) mating types. Besides the mean we also plot the 95%-confidence intervals for each initial condition. The mutation rate was chosen as 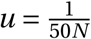 i.e. one mutation every 50 generations on average. Further details on the simulation can be found in the SI. The heuristic prediction *N*^1/3^ works well for lower population sizes but underestimates larger population sizes where the system becomes very stable due to the larger number of individuals of each mating type in equilibrium. In this regime the heuristic argument that extinctions can be described by fluctuations around the stationary value is violated. On the other hand, the formal prediction (red line) follows the increase observed in the data. For large population sizes the computation of this prediction does not give results. Therefore it ends at around a population size of 8000. The dashed lines are bounds on the number of mating types obtained in (Constable and Kokko, 2018) (their equation (3)) which envelope our prediction. (b) Varying the mutation rate *u = N*^−*β*^ affects the predicted number of mating types. While for low mutation rates the number of mating types is well approximated by our prediction, for higher mutation rates (more than one mutation per generation) we see that the number of mating types increases rapidly and is not covered by our the *N*^1/3^-prediction while the rigorous estimate still captures the simulated data. The initial value number of mating types in the simulations is set to 10.

In order to assess the validity of this result we also derived a theoretically rigorous estimate. The method again relies on the analysis of the dynamics of a focal mating type, i.e. equation (2.3), see also (Wright, 1939) for the methodology. However, instead of comparing the single components of the equation we explicitly compute the stationary distribution *f* of this equation, a Gaussian. Its values *f* (*k*/*N*) represent the probabilities of observing *k* individuals carrying the focal mating type in the total population. We then identify the extinction rate of a mating type as the product of *f* (1/*N*), the probability of having exactly one individual of that mating type, and the death rate of this individual 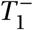, see equation (2.2). The “birth rate” of novel mating types is given by the product of mutation rate *u* and the overall reproduction rate of the population (cf. SI, Section 2.2 for details). Comparing these two quantities for a given population size *N* and a certain number of mating types *R* yields

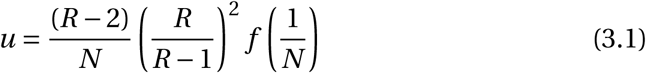

as the mutation rate necessary to maintain *R* mating types in that population.

We find that for low mutation rates and intermediate population sizes the heuristic and rigorous estimate are very similar, see the corresponding lines in Figure 4a and SI, Section 2.3. Intermediate refers to the population sizes where Brownian deviations from the stationary distribution will result in a mating type extinction. This intuition is violated for large population sizes where a mating type is carried by several hundreds (or even thousands) of individuals, meaning that deviations from stationarity do not imply the extinction of a certain mating type allele.

### 3.2 Invasion of new mating types

New mutants, as the sole representatives of their mating type, are subject to both the strongest selection and the greatest risk of being lost by drift. So what is the probability that a novel mating type becomes established in a population? If the rare mutant can avoid immediate extinction, negative frequency-dependence will drive it to higher and higher frequencies, and the risk of extinction will decrease until the mutant reaches a high enough frequency that its survival is guaranteed by the deterministic dynamics (Figure 5a). A small increase in frequency causes a large drop in the risk of extinction but has little effect on the likelihood of encountering an incompatible mate. Even in relatively small populations the risk of extinction will become negligible long before the frequency of the mutant is high enough that the risk of encountering an incompatible mating partner becomes problematic; the selective advantage at these low frequencies (i.e. below the stochasticity threshold) remains essentially constant.

**Figure 5:**
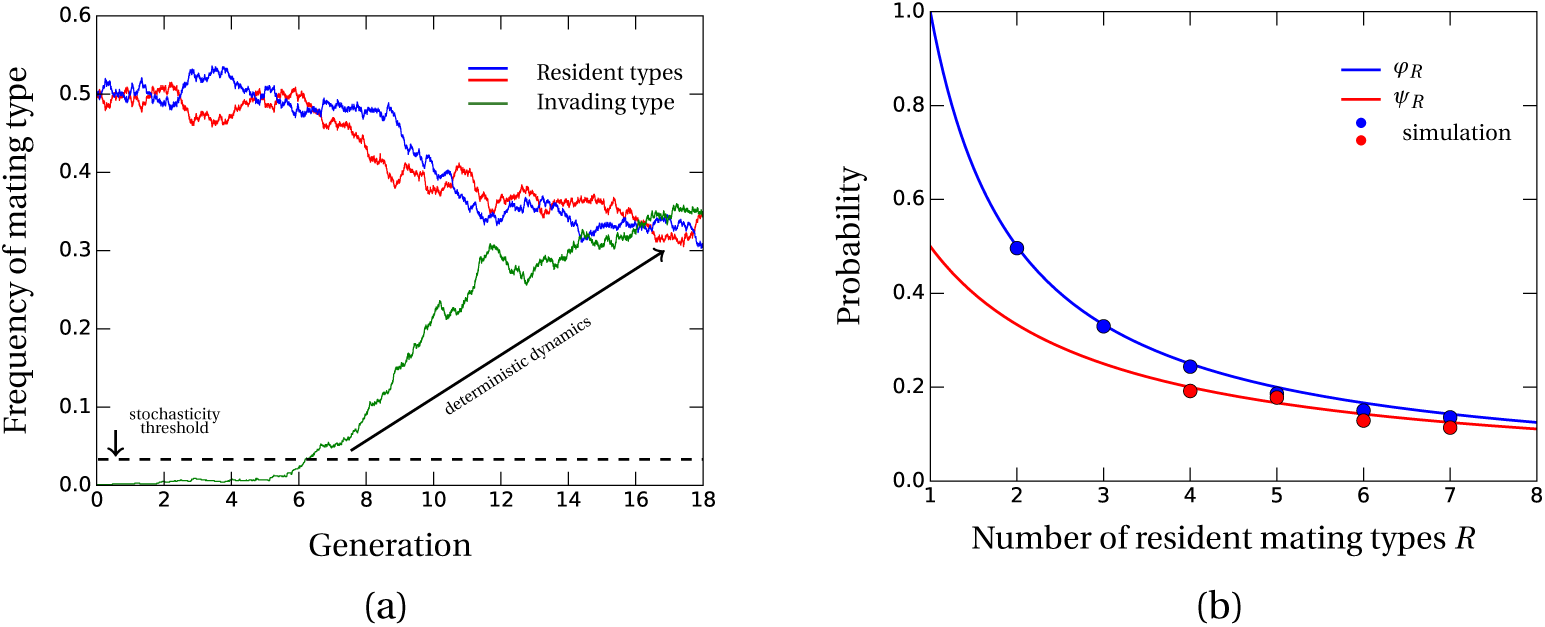
New mating types can invade if they overcome the stochasticity threshold. (a) A typical invasion behavior of a new mating type is plotted. While in low frequency the invading type is prone to stochastic fluctuations, eventually causing extinction. For higher frequencies (above the stochasticity threshold) the deterministic dynamics carry the new mating type towards the new deterministic equilibrium at 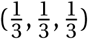. (b) Choosing the population size as *N =* 250 (larger values improve the fit), we plot our predictions for the invasion (blue) and turnover (red) probability and compare them with simulation results. Invasion data points are derived from 10,000 runs with the resident population being close to its equilibrium 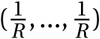 and one single new mating type individual. The same initial state is assumed for the 1,000 turnover simulations. Simulation results for *R =* 2, 3 are missing in case of the turnover probability since populations consisting of that few mating types are too stable for a mating type to go extinct. This is consistent with our prediction that below 250^1/3^ ≈ 6 deterministic dynamics outweigh stochastic drift meaning extinctions are very rare.

Computing the establishment probability therefore means calculating the survival probability of the rare mating type at frequencies small enough that the risk of encountering an incompatible mate can be ignored. Given *R* different mating types in the population, we find that the probability of establishment of the (*R* + 1)-th type is (the detailed derivation is given in SI, Section 3)

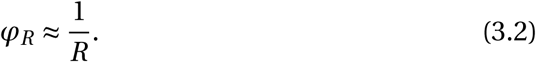

The inverse relationship between the establishment probability *φ* and the number of resident mating types can be explained by the selective advantage of a rare mutant when compared to a resident type. While a rare type will successfully reproduce with probability 1, a resident mating type has only probability 1 − 1/*R* resulting in a fitness difference of 1/*R*. Thus, when the number of resident types is low, 1/*R* is large and thus, the extinction risk of the mutant is small. Contrarily, when the number of resident mating types is high, i.e. 1/*R* becomes very small, the reproductive advantage of the mutant becomes smaller and hence its extinction risk increases. Additionally, equation (3.2) implies that for very large numbers of resident mating types the invasion probability is essentially zero resulting in no novel mating types establishing in the population.

Figure 5b shows a comparison of equation (3.2) with stochastic simulations in a population with *N =* 250 individuals. The accuracy of our prediction is limited to the region where deterministic dynamics dominate the system. Otherwise the intuition that frequency-dependent selection stabilizes the rare mutant fails and no analytical approximation is available. Hence, larger population sizes *N* not only improve the quality of our approximation but also increase the range for *R* where our estimate is applicable.

### 3.3 The turnover probability of mating types

We have shown that finite populations will eventually reach a relatively stable number of mating types. However, the stability of the number of mating types does not imply evolutionary stasis. Resident mating types can be continuously replaced by new mating types in a process referred to as the “turnover of the sexes” (Figure 6; see also (Iwasa and Sasaki, 1987)).

**Figure 6:**
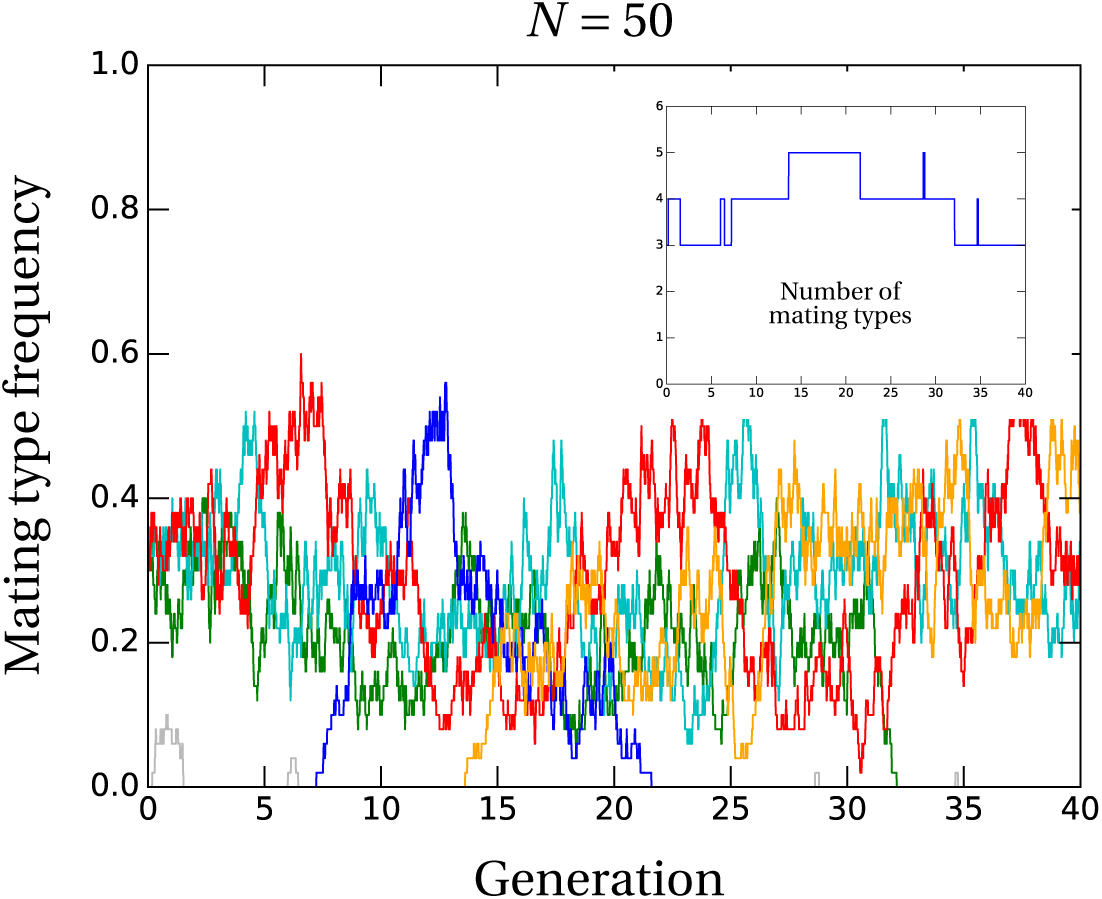
The number of types in a population remains stable but mating types are rapidly lost and replaced in small populations. The process “turnover of sexes” in a population of size *N =* 50. Initially, three mating types are present at equal frequencies (red, green, cyan). After two unsuccessful invasions (grey) a new mating type invades the resident population (blue). This invasion is followed by a second invasion (orange) before an established mating type is lost (blue). Due to an excess of mating types eventually also the green mating type is lost from the population resulting in a new population consisting of the orange, red and cyan type. The inset shows the number of mating types present in the population over time.

Computation of the probability for such a turnover to happen is straightforward in our setting. Since it can be seen as a two step process, invasion of a mutant type followed by extinction of a resident mating type, the probability is given by

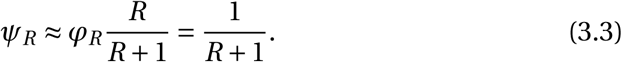

The invasion probability as calculated in the previous section in equation (3.2) is 1/*R* while the extinction probability of a resident type is given by *R*/(*R* + 1) (all mating types behave equally).

As demonstrated in Figure 5b our estimate of the turnover probability fits the data points derived from stochastic simulations. For populations far below the maximal number of mating types (*R =* 2, 3) the population is too stable such that effectively no turnover takes place indicating large extinction times for low resident numbers. If anything, the new mating type establishes and by that increases the overall number of mating types in the population. This explains the missing data points for *R =* 2, 3. Once again our estimate improves for larger values of *N*, similar to the estimate of the invasion probability. The simulations are restricted to *N =* 250 since for larger population sizes extinction times of a mating type exceed the computational time constraints.

## 4. Discussion

Even though rare mating types have a significant selective advantage in a sexually reproducing organism with self-incompatible gametes, most species have a very small number of mating types - typically only two (James, 2015). Here, we present an analytical explanation for the discrepancy between the low number of types observed in natural populations and the deterministic prediction. Heuristically identifying the point at which the deterministic dynamics – which maintain all types at equal frequency – are balanced by the stochastic dynamics – which can cause rare types to be lost from the population, we find that the maximum number of mating types that can be supported by a finite population is of order *N*^1/3^ which falls within previously identified bounds (Constable and Kokko, 2018). We show that this value is a good approximation for small to intermediate population sizes using a rigorous numerical evaluation adapted from models developed for investigating gametophytic self-incompatibility in plants (Wright, 1939; Fisher, 1958; Wright, 1960, 1964; Ewens, 1964; Nagylaki, 1975; Yokoyama and Hetherington, 1982; Slatkin and Muirhead, 1999), reviewed in (Clark and Kao, 1994). Thus, accounting for both deterministic negative frequency-dependence and genetic drift reveals that the number of mating types in real populations is far below the naïve deterministic expectation (Billiard et al., 2011; Constable and Kokko, 2018).

Still, when considering biological reality our model only accounts for the competing influences of mutation, negative frequency dependent selection, and genetic drift. The number of mating types supported by the balance of these forces should therefore be considered a maximum. Many other factors, not included in our model, could drive the number of types in a population below this limit. Chief among these factors is that our model assumes that newly arising mating types are fully compatible with all resident types. This could be unrealistic if resident types have co-evolved strong interactions. New mutants suffering reduced compatibility will be less likely to invade, limiting the number of types in a population to be below our calculated maximum (Power, 1976; Hadjivasiliou and Pomiankowski, 2016). Indeed, in systems where compatibility is determined by highly specific ligand receptor interactions, it seems unlikely that a random mutant would not experience reduced mating efficiency (Hoekstra, 1987). Nevertheless, pheromone-receptor mate recognition systems involving multiple ligands do exist. The ciliate *Euplotes raikovi* can at least partially distinguish between self-pheromones and up to eight different non-self pheromones (Luporini et al., 2016). Nieuwenhuis et al. (2013) suggested that this problem might be reduced in tetrapolar systems (mating types are defined by two loci) where pheromone-receptor compatibility is uncoupled from heterodimerizing homeodomain proteins that regulate the expression of mating types, allowing both loci to evolve independently. Indeed, multiallelic haploid incompatibility found in certain bipolar fungi may have evolved in their tetrapolar ancestors (James et al., 2006). Of course, the number of possible combinations of compatible homeodomain proteins is almost certainly limited as well (Perrin, 2012).

Our model is based on the life cycle of *S. cerevisiae*, but for simplicity we have only examined sexual reproduction. These yeast are actually facultatively sexual, and alternate between sexual and asexual models of reproduction, although the relative frequencies of these two modes remains highly controversial (Kelly and Wickner, 2013). As reported by (Constable and Kokko, 2018) populations with a high frequency of asexual reproduction are expected to carry fewer mating types than equivalent sexual populations. By decreasing the contribution of sexual reproduction to fitness, asexuality effectively weakens the deterministic dynamics relative to the stochastic dynamics, shifting the balance between these two forces to a lower number of mating types. *S. cerevisiae* is also capable of switching between mating types when compatible partners are rare. Previous simulation studies found that mating type switching is mostly favored when allele frequencies, due to low population sizes or high mating type numbers, are prone to strong fluctuations so that the most common type frequently fails to find a compatible partner (Paixão et al., 2011; Hadjivasiliou et al., 2016). The exact consequences of switching on the number of mating types that can be maintained in a population remains to be explored. Further issues that might limit the number of mating types are reviewed by (Billiard et al., 2011).

Comparing our results to empirical observations is further hindered by difficulties in accurately estimating effective population sizes, particularly in microbes. Estimates of the effective population sizes of a single species can vary by several orders of magnitude (Katz et al., 2005) but some unicellular protists are thought to have very small effective population sizes: temporal estimates from a single geographical location of the marine dinoflagellate *Pentapharsodinium dalei* revealed an effective population size in the order of 100 (Watts et al., 2013).

In large populations, or in populations with low mutations rates, we show that the number of mating types will be stable. However, this should not be confused with evolutionary stasis. As originally suggested by (Iwasa and Sasaki, 1987), the identity of mating types present in a population may change over time. As a first step towards an analytical solution for this turnover rate, we derive the probability of invasion of a rare mating type. It is given as the inverse of the number of resident mating types. This can be interpreted as the fitness advantage of a novel mutant when compared to a resident mating type: low resident numbers yield large selective benefits to the rare mutant while large resident numbers lower the fitness of a mutant. Together with the mutation rate this invasion probability gives the establishment rate of newly arising mating types. Still, to predict the turnover rate one would additionally need the extinction time of a resident mating allele. Previous studies in the context of the major histocompatibility complex and of plant self-incompatibility systems have estimated the extinction time of alleles under constant balancing selection (Takahata, 1990; Vekemans and Slatkin, 1994). An extension and application of these techniques to our model is beyond the scope of this study but see (Czuppon and Constable, 2019) for an assessment of extinction times in the here studied setting.

In conclusion, our findings add new theoretical results and mechanistic insights on the evolutionary dynamics of mating type alleles in a haploid self-incompatibility system. Investigations of the evolution of multiallelic loci subject to negative frequency dependent selection have been studied extensively. Applying these previously developed methods to our Moran-model of haploid self-incompatibility we provide two estimates on the maximum number of alleles in a finite population. The simplicity of the heuristic derivation has the advantage of obtaining an explicit prediction of the number of mating types while coming at the cost of being restricted to low mutation rates and intermediate population sizes. The formal analysis on the other hand, provides a more robust estimate for the mutation-selection-drift balance while being reliant on numerical evaluation since no closed form solution is attainable. Moreover, we compute invasion and turnover probabilities of newly arising mating types which, to the best of our knowledge, have not been studied for any highly polymorphic system before. This latter approach can be extended to the investigation of alleles conferring differential fitness, allowing the study of more complex mating systems, meiotic drive, or other multiallelic systems subject to balancing selection including certain gamete recognition proteins (Tomaiuolo and Levitan, 2010).

## Notes

#### Summary of Updates

new version

