## Supplementary Information for "Evolution of mating types in finite populations: the precarious advantage of being rare"

#### 1 Derivation of equation (2.3) in the main text

In this section we derive the stochastic differential equation (2.3) from the main text that describes the dynamics of a focal mating type. For that reason, we will explicitly calculate the infinitesimal change in the expectation, the variance and covariances of the process which translate to the deterministic drift and stochastic diffusion term of the stochastic differential equation. For a more detailed description of these quantities and their relation we refer to (Etheridge, 2012, Chapter 3).

Let us recall the transition rates for type $M_i$ individuals. We denote the absolute number of $M_i$ individuals by $X_i$ and the corresponding frequency by $x_i = X_i/N$. As explained in the main text, the rate to go from $k$ type $M_i$ individuals to $k+1$ individuals reads

$$T_k^{i+} = \frac{1}{2}k\frac{N-k}{N-1}\frac{N-k-1}{N-2} + \frac{1}{2}(N-k)\frac{k}{N-1}\frac{N-k-1}{N-2} = k\frac{N-k}{N-1}\frac{N-k-1}{N-2}, \quad (1.1)$$

whereas the rate of decrease is given by

$$T_k^{i-} = \sum_{j\neq i} X_j \frac{N - X_j - k}{N-1}\frac{k}{N-2} + \sum_{j\neq i} X_j \frac{k}{N-1}\frac{k-1}{N-2} = \sum_{j\neq i} X_j \frac{N - X_j - 1}{N-1}\frac{k}{N-2}. \tag{1.2}$$

This yields the infinitesimal expected change, i.e. the change in a small time interval $\delta t$, in relative numbers $x_i$

$$\begin{aligned}
\mathbb{E}[x_i(t+\delta t) - x_i(t)|x(t)] &= \frac{\delta t}{N}\left(T^{i+}_{x_i(t)N} - T^{i-}_{x_i(t)N}\right) \\
&\approx \delta t \left[x_i(t)(1-x_i(t))^2 - \sum_{j\neq i} x_j(t)(1-x_j(t))x_i(t)\right] \\
&= \delta t \left[x_i(t)\left((1-x_i(t))^2 - \sum_{j\neq i} x_j(t)(1-x_j(t))\right)\right] \\
&= \delta t \left[x_i(t)\left(1-x_i(t) - \sum_{j=1}^{R} x_j(t)(1-x_j(t))\right)\right] \\
&= \delta t \left[x_i(t)\left(\sum_{j=1}^{R} x_j(t)^2 - x_i(t)\right)\right],
\end{aligned} \tag{1.3}$$

where we have set $R$ as the number of present mating types.

Next, we proceed deriving the change in variance in a short time interval. Neglecting terms of order $O(\delta t^2)$ we find (for $\Delta x_i(t+\delta t) = x_i(t+\delta t) - x_i(t)$)

$$\begin{aligned}
Var[\Delta x_i(t+\delta t)|x(t)] &= \mathbb{E}[(x_i(t+\delta t) - x_i(t))^2|x(t)] - \mathbb{E}[x_i(t+\delta t) - x_i(t)|x(t)]^2 \\
&= \mathbb{E}[(x_i(t+\delta t) - x_i(t))^2|x(t)] - O(\delta t^2) \\
&\approx \frac{1}{N^2}\delta t\left(T^{i+}_{x_i(t)N} + T^{i-}_{x_i(t)N}\right) \\
&\approx \frac{\delta t}{N}\left[x_i\left((1-x_i(t))^2 + \sum_{j\neq i} x_j(t)(1-x_j(t))\right)\right] \\
&= \frac{\delta t}{N}\left[x_i\left(2 - 3x_i + 2x_i^2 - \sum_{j=1}^{R} x_j^2\right)\right].
\end{aligned} \tag{1.4}$$

Lastly, for the covariances we find (again neglecting terms of order $(\delta t)^2$)

$$\begin{aligned} Cov[\Delta x_i, \Delta x_j | x(t)] &= \mathbb{E}[(\Delta x_i - \mathbb{E}[\Delta x_i])(\Delta x_j - \mathbb{E}[\Delta x_j]) | x(t)] \\ &= \mathbb{E}[\Delta x_i \Delta x_j | x(t)] - \mathbb{E}[\Delta x_i | x(t)]\mathbb{E}[\Delta x_j | x(t)] \\ &\approx -\frac{\delta t}{N}\left(x_i(1-x_i)x_j + x_j(1-x_j)x_i\right) \\ &= -\frac{\delta t}{N} x_i x_j (2 - x_i - x_j). \end{aligned} \tag{1.5}$$

Collecting all the terms we find the following stochastic diffusion process describing the dynamics of a focal mating type :

$$\begin{aligned} dx_i &= x_i\left(\sum_j x_j^2 - x_i\right)dt - \sum_{j\neq i}\sqrt{\frac{x_i x_j(2-x_i-x_j)}{N}}dW_j \\ &\qquad + \sqrt{\frac{x_i\left(2-3x_i+2x_i^2-\sum_j x_j^2\right)}{N}}dW_i \\ &= x_i\left(\sum_j x_j^2 - x_i\right)dt - \sum_{j\neq i}\sqrt{\frac{C_{ij}(x)}{N}}dW_j + \sqrt{\frac{\sigma_i(x)}{N}}dW_i, \end{aligned} \tag{1.6}$$

where $C_{ij}(x) = x_i x_j(2 - x_i - x_j)$ and $\sigma_i(x) = x_i\left(2-3x_i+2x_i^2-\sum_j x_j^2\right)$ are the variables used in the main text and $(W_i)_{i=1,\ldots,R}$ are independent one-dimensional Brownian motions. The first part of the equation, the expected change in the mean, reflects the deterministic evolution of the system. The stochasticity on the other hand is captured by the Brownian motions which correspond to the infinitesimal change of the variance and covariances, all with respect to the focal mating type.

#### 1.1 Including mutations

In a next step we include mutations, the mechanism which generates new mating types. We implement this during the reproduction step meaning that there is a probability that after a successful mating the offspring expresses a completely new mating type. More precisely, we also exclude mutations to already existing mating types, thus assuming a variant of the infinitely many alleles model. Denoting by $u$ the mutation rate, the increase rate of a focal mating type $M_i$ transforms to

$$U_k^{i+} = (1-u)\left(k\frac{N-k}{N-1}\frac{N-k-1}{N-2}\right).$$

Additionally, we have an increase rate due to mutations if a mating type $M_i$ is not present in the population, i.e.

$$U_0^{i+} = \frac{u}{N-R}\sum_{j=1}^{R} X_j \frac{(N-X_j)}{N-1}, \tag{1.7}$$

where the denominator $N-R$ represents the fact that each non-present mating type is equally likely to be obtained through a mutation with $N$ being the maximal number of possible mating types in a population of $N$ individuals.

Finally, the decrease rate now has an additional mutation term such that it can be written as

$$U_k^{i-} = \sum_{j\neq i} X_j \frac{N-X_j-1}{N-1}\frac{k}{N-2} + u\frac{k}{2}\frac{N-k}{N-1}\frac{k-1}{N-2}.$$

These are the rates used in the individual based simulations. A derivation of a stochastic diffusion equation including mutations is not necessary since we will base all of our analysis below on the case of weak mutations in which case the corresponding limit reduces to the one derived in equation (1.6) above.

### 2 The expected number of mating types

#### 2.1 Heuristic estimate

Let us first derive the heuristic estimate of $N^{1/3}$. We start by considering the deterministic component of equation (1.6) in case $x_i > 0$, i.e.

$$x_i\left(\sum_{j=1}^{R} x_j^2 - x_i\right).$$

We assume that the population is in its internal equilibrium with $R$ mating types, i.e. at $(1/R,...,1/R)$. Writing $R$ in terms of $N$ we set $R = N^{\alpha}$ for $\alpha > 0$ and thus, the frequency of a single mating type can be written as

$$x_i = \frac{1}{R} = N^{-\alpha}, \quad \text{for all } i.$$

Now the order of the deterministic component can be written as

$$x_i\left(\sum_{j=1}^{R} x_j^2 - x_i\right) \sim N^{-\alpha}\left(N^{\alpha}N^{-2\alpha} - N^{-\alpha}\right) \sim N^{-2\alpha}. \tag{2.1}$$

Hence, the deterministic dynamics are of order $N^{-2\alpha}$. For the stochastic counterpart of equation (1.6) we recall the functions $C_{ij}(x)$ and $\sigma_i(x)$. They read as

$$C_{ij}(x) = x_i x_j(2 - x_i - x_j) \quad \text{and} \quad \sigma_i(x) = x_i\left(2 - 3x_i + 2x_i^2 - \sum_{j=1}^{R} x_j^2\right).$$

Therefore, the order of the stochastic fluctuations is given by

$$C_{ij}(x) \sim N^{-2\alpha} \quad \text{and} \quad \sigma_i(x) \sim N^{-\alpha}.$$

Due to the additivity of independent Brownian motions we find

$$\sqrt{\sum_{j\neq i}\frac{C_{ij}(x)}{N} + \frac{\sigma_i(x)}{N}} \sim \left(N^{\alpha}N^{-(2\alpha+1)} + N^{-(\alpha+1)}\right)^{1/2} \sim N^{-\frac{\alpha+1}{2}} \tag{2.2}$$

for the order of the stochastic dynamics. Solving (2.1) > (2.2) which means that the deterministic dynamics dominate the system resulting in extinctions to be unlikely, gives

$$N^{-2\alpha} > N^{-\frac{\alpha+1}{2}} \quad \Leftrightarrow \quad \frac{1}{3} > \alpha. \tag{2.3}$$

This states that in a population of $N$ individuals with a sufficiently low mutation rate $u$ and intermediate values of $N$ we would expect around $N^{1/3}$ different mating types, the result stated in the main text.

#### 2.2 Rigorous estimate

In this section we derive an estimate for the number of mating types following the methods developed by Wright in the context of self-incompatibility systems in plants (Wright, 1939, 1960, 1964). The idea is to use the stationary distribution $f$ of a focal mating type, more precisely the value of this distribution at the extinction boundary with one sole individual, i.e. $f(1/N)$. Then comparing the decrease rate from this terminal class with the "birth"-rate of new mating type alleles in equation (1.7) will give an estimate on the number of mating types.

For the estimation of the stationary distribution of equation (2.3) in the main text we will make use of the general form as stated for example in (Ewens, 2004, Chapter 4.5) which, when neglecting the covariance terms, reads

$$f(x) = \frac{c}{\sigma^2(x)} \exp\left(2\int_0^x \frac{\mu(y)}{\sigma^2(y)} dy\right),$$

where $c$ is the normalization constant, $\mu$ the deterministic and $\sigma^2$ the variance term.

Using the diffusion approximation from equation (1.6) (assuming weak mutation) and fixing the non-$M_i$ mating types as a point measure to $1/R$, i.e. $\sum_{j=1}^R x_j^2 = (R-1)/R^2 + x_i^2$ we find

$$\begin{aligned}\mu(x) &= x\left(\frac{1}{R}\left(1-\frac{1}{R}\right) - x(1-x)\right),\\ \sigma^2(x) &= \frac{x}{N}\left(2-3x+x^2-\frac{1}{R}\left(1-\frac{1}{R}\right)\right) \approx \frac{x}{N}\left(2-3x-\frac{1}{R}\right),\end{aligned}$$

where the approximation is based on $x^3/N$ and $x/(R^2N)$ being of negligible order of magnitude.

Using these values we can solve the integral in the stationary distribution with symbolic programming languages (e.g. *Mathematica*) which yields

$$\int_0^x \frac{\mu(y)}{\sigma^2(y)} dy \approx \frac{N}{54}\left[3x\left(2+\frac{2}{R}-3x\right)+4\left(1-\frac{2}{R}\right)^2 \log\left(2-\frac{1}{R}-3x\right)\right].$$

We note, that the logarithm is well-defined since for $R=2$ the prefactor vanishes and otherwise the condition $2-1/R-3x>0$ holds since $x \le 1/2$, a reasonable assumption given that we assume the other (at least) two mating types are at frequency $1/3$ and $N$ is sufficiently large.

Putting things together we find the following expression for the stationary distribution

$$f(x) \approx \frac{Nc}{x}\left(2-3x-\frac{1}{R}\right)^{\frac{4N}{27}\left(1-\frac{2}{R}\right)^2-1} \exp\left(\frac{Nx}{9}\left(2-3x+\frac{2}{R}\right)\right). \tag{2.4}$$

Evaluating at $x = 1/N$ gives

$$f\left(\frac{1}{N}\right) \approx N^2 c\left(2-\frac{3}{N}-\frac{1}{R}\right)^{\frac{4N}{27}\left(1-\frac{2}{R}\right)^2-1} \exp\left(\frac{1}{9}\left(2-\frac{3}{N}+\frac{2}{R}\right)\right).$$

The constant $c$ can be computed numerically by summing over all possible states (assuming that states larger than $1/2$ are not relevant) so that we are left with determining the decrease rate when we have one individual present in the population. In fact, having one individual of a certain mating type, we can assume that the rest of the population is close to the (new) steady state, i.e. $x_j \approx 1/(R-1)$. Therefore, using equation (1.2) (scaled by $1/N$) we find for the extinction rate

$$\begin{aligned}\mu_R &= T_1^- f\left(\frac{1}{N}\right) \approx \frac{1}{N}\left(1-\frac{1}{R-1}\right) f\left(\frac{1}{N}\right) = \frac{(R-2)}{N(R-1)} f\left(\frac{1}{N}\right) \\ &\approx Nc\left(\frac{R-2}{R-1}\right)\left(2-\frac{3}{N}-\frac{1}{R}\right)^{\frac{4N}{27}\left(1-\frac{2}{R}\right)^2-1} \exp\left(\frac{1}{9}\left(2-\frac{3}{N}+\frac{2}{R}\right)\right) .\end{aligned} \tag{2.5}$$

We note that the last expression in the first line of the previous formula is similar to the corresponding equation obtained by Wright in his analysis, compare for example (Wright, 1939, p. 541).

We will need to compare $R\mu_R$ (each of the mating types can die out) with the introduction of new mating types, i.e. equation (1.7). Again scaling that equation we find as an increase rate for the number of mating types when assuming that the present mating types are in equilibrium,

$$\lambda_R = \frac{u}{(N-R)}\left(1-\frac{1}{R}\right) = \frac{u}{(N-R)R}(R-1) . \tag{2.6}$$

Multiplying by the number of possible new mating types given by $N-R$ (there can be at most $N$ mating types in a population of $N$ individuals) we find

$$(N-R)\lambda_R = R\mu_R \quad \Leftrightarrow \quad u = \frac{(R-2)}{N}\left(\frac{R}{R-1}\right)^2 f\left(\frac{1}{N}\right) . \tag{2.7}$$

#### 2.3 Comparison between heuristic and rigorous prediction

In the following we explore the population size parameter to evaluate in which regimes the heuristic – $N^{1/3}$ – prediction is a good approximation for the number of self-incompatible mating types. Comparing this to the rigorous prediction from equation (2.7) shows that up until approximately 3000 individuals in a population both solutions agree, see also Figure 1. For larger population sizes the assumptions of the $N^{1/3}$-derivation are not valid anymore. More precisely, the number of individuals carrying a certain mating type allele in stationarity

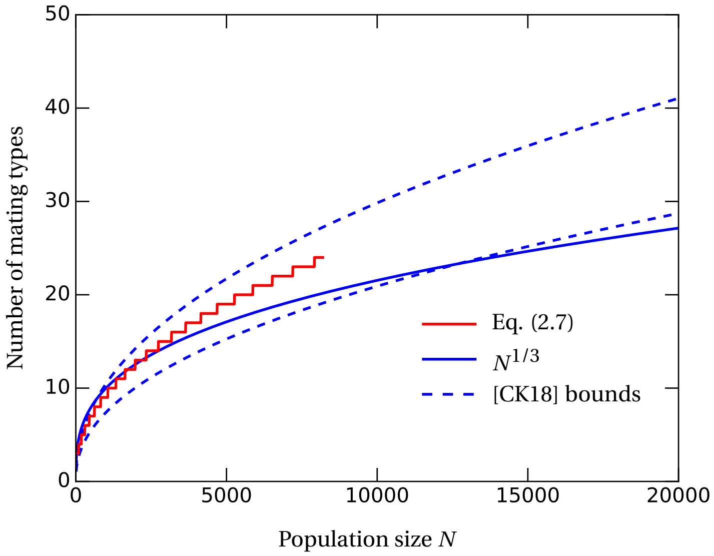


Figure 1: **Comparing the heuristic (blue) and rigorous (red) prediction reveals the population sizes where the heuristic approximation is valid.** While the two estimates agree for small population sizes they diverge for larger population sizes. The rigorous analysis seems to remain within the bounds obtained in (Constable and Kokko, 2018) while the $N^{1/3}$ prediction will cross the lower bound for very large population sizes (around $N = 13,000$).

is too high for a Brownian stochastic fluctuation to cause an extinction event. The same reasoning explains why for very large population sizes ($\sim 13000$) the $N^{1/3}$ prediction will leave the predicted boundaries obtained in (Constable and Kokko, 2018).

### 3 Invasion Probability

In order to calculate the probability for a new mating type to establish itself in the population we employ techniques from branching process theory in a constant environment. This is reasonable since we only need to compute the probability for the new mating type to reach a certain threshold frequency from where deterministic dynamics take over and pull the frequency up to the stable internal equilibrium. However, this is only reasonable in the regime in which the deterministic dynamics dominate the stochastic fluctuations, i.e. when the number of mating types is smaller than $N^{1/3}$ - see Section 3.1 in the main text. Hence, let us assume that the number of mating types in the population is $R < N^{1/3}$. A new mating type arises and we can compute its birth and death rates under the assumption that the other $K$ mating types are close to their equilibrium frequencies, i.e. $\frac{1}{R}$. Then we find

$$T_k^+ = k\frac{N-k}{N-1}\frac{N-k-1}{N-2} \approx k,$$

since we assume $N$ to be large and $k$, the number of individuals having the new mating type, to be negligibly small, i.e. $\frac{N-k}{N} \approx 1$. This is equivalent to the assumption that the individuals of the novel mating type evolve independently of each other, the basic assumption of a branching process. For the death rate of individuals expressing the new mating type we have

$$T_k^- = \sum_{i=1}^{R} k_i \frac{N-k_i-1}{N-1}\frac{k}{N-2} \approx kR\frac{1}{R}\left(1-\frac{1}{R}\right) = k\left(1-\frac{1}{R}\right).$$

Note, that this approximation also holds if there is more than one rare mutant mating type in the population. One of those new mating types will eventually reach a critical threshold from where deterministic dynamics will bring it to the new internal equilibrium $\frac{1}{R+1}$. However, a rigorous analysis of this situation is beyond the scope of this work. Hence, we restrict ourselves to low mutation rates which allow us to work in a situation where we only have one invading mating type at a time. Then the above transition rates hold until the new mating type hits a low threshold value from where on its survival is guaranteed due to the deterministic dynamics, see also Figure 5a in the main text. To calculate the hitting probability of this (arbitrarily set) threshold we use branching process theory, i.e. we calculate the survival probability of this auxiliary process. Since our system allows for a stable fixed point in the deterministic system the new mating type will, after crossing the threshold, increase the number of mating types.

Let us now proceed computing this invasion or establishment probability which we call $\varphi_R$. As we will see, it is solely dependent on the present number of mating types $R$. Noting, that the number of individuals having the invading mating type is a continuous-time birth-death process we can apply known theorems for computing the survival probability of it, see for instance (Allen,

2011, Theorem 6.2). Thus, the extinction probability for a birth-death process with one initial individual is given by the formula

$$p_{\text{ext}} = \frac{\sum_{k=1}^{\infty} \frac{T_1^- \cdots T_k^-}{T_1^+ \cdots T_k^+}}{1 + \sum_{k=1}^{\infty} \frac{T_1^- \cdots T_k^-}{T_1^+ \cdots T_k^+}}. \tag{3.1}$$

Inserting our rates from above and applying the formula for the geometric series this yields

$$\begin{aligned} \varphi_R = 1 - p_{\text{ext}} &= 1 - \frac{\sum_{k=1}^{\infty} \left(1 - \frac{1}{R}\right)^k}{1 + \sum_{k=1}^{\infty} \left(1 - \frac{1}{R}\right)^k} \\ &= 1 - \frac{1}{\frac{1}{1-\left(1-\frac{1}{R}\right)}} \left(\frac{1}{1 - \left(1 - \frac{1}{R}\right)} - 1\right) \\ &= 1 - \frac{1}{R}(R-1) = \frac{1}{R}. \end{aligned} \tag{3.2}$$

### 4 Simulations

The data used for the estimates in Figure 4 were simulated in the following way. One simulation starting either with 3, 10 or 20 mating types in the corresponding deterministic equilibrium was first given 20,000 generations to equilibrate. Then we sampled the number of mating types every generation, i.e. after $N$ transitions, for in total 1,000 generations. Each initial condition was run for 50 independent runs such that each symbol is an average of 50,000 data points.
